# Identification of functional white matter networks in BOLD fMRI

**DOI:** 10.1101/2023.09.08.556881

**Authors:** Alexa L. Eby, Lucas W. Remedios, Michael E. Kim, Muwei Li, Yurui Gao, John C. Gore, Kurt G. Schilling, Bennett A. Landman

## Abstract

White matter signals in resting state blood oxygen level dependent functional magnetic resonance (BOLD-fMRI) have been largely discounted, yet there is growing evidence that these signals are indicative of brain activity. Understanding how these white matter signals capture function can provide insight into brain physiology. Moreover, functional signals could potentially be used as early markers for neurological changes, such as in Alzheimer’s Disease. To investigate white matter brain networks, we leveraged the OASIS-3 dataset to extract white matter signals from resting state BOLD-FMRI data on 711 subjects. The imaging was longitudinal with a total of 2,026 images. Hierarchical clustering was performed to investigate clusters of voxel-level correlations on the timeseries data. The stability of clusters was measured with the average Dice coefficients on two different cross fold validations. The first validated the stability between scans, and the second validated the stability between subject populations. Functional clusters at hierarchical levels 4, 9, 13, 18, and 24 had local maximum stability, suggesting better clustered white matter. In comparison with JHU-DTI-SS Type-I Atlas defined regions, clusters at lower hierarchical levels identified well defined anatomical lobes. At higher hierarchical levels, functional clusters mapped motor and memory functional regions, identifying 50.00%, 20.00%, 27.27%, and 35.14% of the frontal, occipital, parietal, and temporal lobe regions respectively.

## 1. INTRODUCTION

Blood oxygen level dependent functional magnetic resonance imaging (BOLD-fMRI) is a common noninvasive technique used to measure neuronal activity through the spontaneous hemodynamic responses in blood^1,2^. At resting state, BOLD-fMRI is used to examine the baseline functional signaling within the brain during which the subject is not completing a specific task. Using these signals, grey matter has been used extensively to create functional maps in the brain^3^. This grey matter functional mapping is important for understanding aging effects and diseases, such as Alzheimer’s Disease^4,5^ In contrast to grey matter, white matter is less voluminous and has smaller BOLD signals especially at resting state^6,7^. As a result, white matter BOLD signals have previously been discounted or considered noise, but there is growing evidence these white matter signals can provide insight into functional brain regions^7^.

Recent literature has begun investigating the significance of white matter signals in the resting state BOLD-fMRI. The hemodynamic responses of white matter and grey matter have been found to be similar during task and resting state fMRI, providing evidence that white matter signals should not be dicounted^7,8^. Similarly, a recent study has revealed clustered white matter correlations are highly consistent with the identified grey matter correlations^9^ (Figure 1). Furthermore, white matter functional networks have been correlated with grey matter functional networks to investigate the inter-network connectivity^10^. However, this previous work has been completed using k-means clustering techniques only. Our work uses a hierarchical clustering approach to investigate clustering on BOLD-fMRI white matter correlations.

**Figure 1.**
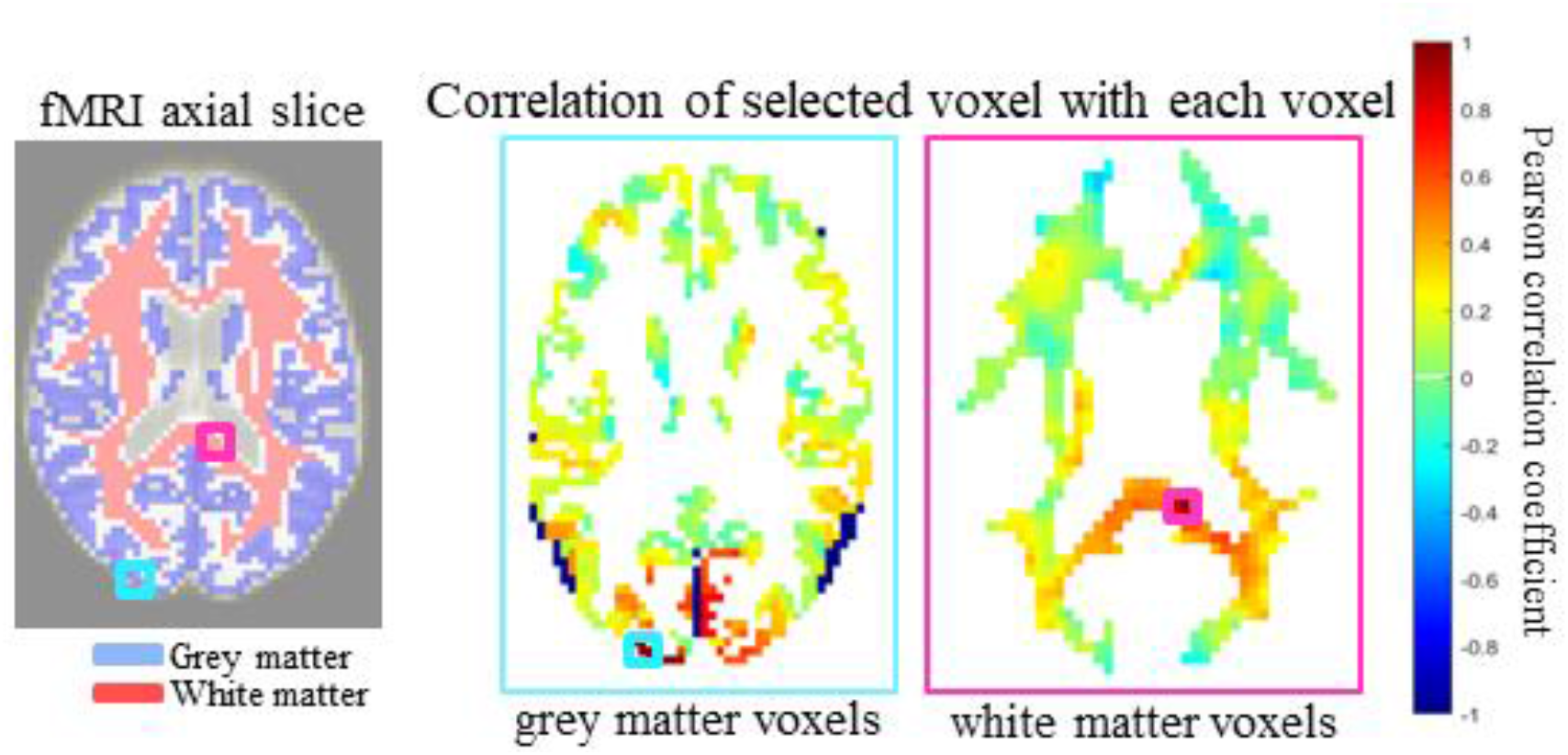
In resting state BOLD fMRI signals there are expected correlations between grey matter voxels which can be used to identify anatomical regions^7^ (left). In a similar manner, we hypothesize correlations can be identified between white matter voxels (right), providing evidence there may be informative white matter correlations in resting state BOLD fMRI data.

**Figure 2.**
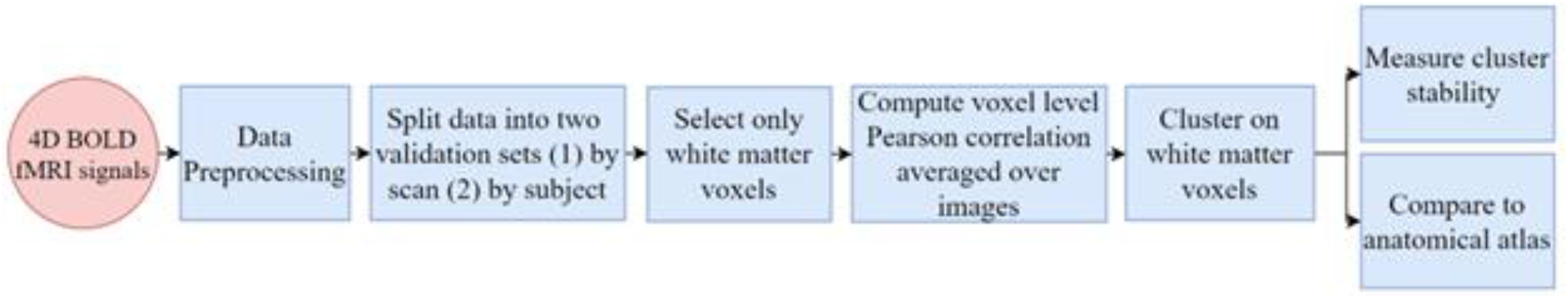
The approach to discover spatial patterns in fMRI imaging using standardized clustering techniques, focusing on white matter voxels. BOLD fMRI signals containing multiple subjects with multiple scans each were preprocessed and standardized using a customized pipeline based on the DPABI toolbox. To measure cluster stability, two validation tests were completed-one set by assigning each scan to a cohort, and the second set by assigning each subject to a cohort. White matter voxels were extracted from the BOLD signals using a group-wise white matter mask. Pearson correlation coefficients were computed for each voxel in the image and averaged across all images. Then, white matter voxels were clustered using average-linkage hierarchical clustering with distance between voxels measured as 1-correlation. Lastly, the average Dice coefficient of clusters at different hierarchical levels was used to identify stable clusters which can be visually compared to know anatomical brain regions. This clustering technique was also compared to the JHU-DTI-SS Type-I Atlas.

## 2. METHODOLOGY

Seven hundred and eleven (cognitively normal, clinical dementia rating = 0, 324 males and 387 females, whose ages ranged between 42 and 95 years) individuals were selected from the OASIS-3 database^11^. Many individuals provide longitudinal data and most of the individuals were scanned twice in a given session, resulting in a total of 2,026 resting state BOLD-fMRI images.

### 2.1 Image Processing

Slice timing and head motion were removed from the fMRI volumes, and then the mean cerebrospinal fluid (CSF) signal and 24 motion-related parameters were modeled as covariates and regressed out from the BOLD signals. The data were then detrended and passed through a temporal filter with a passband frequency of between 0.01 and 0.1 Hz. All these procedures were carried out using a customized pipeline based on the DPABI toolbox^12,13^. The Computational Anatomy Toolbox (CAT12) was then used to segment grey matter, white matter, and CSF tissue based on the T1-weighted images from OASIS-3 data^14^. Using co-registration and normalizing functions in SPM12^15^ the filtered fMRI data, along with corresponding tissue masks, were spatially normalized into MNI space (voxel size = 3 × 3 × 3 mm^3^). As the analyses were restricted to white matter, a group-wise white matter mask was constructed by averaging the white matter parcellations (probability maps) that were derived from CAT12 across all subjects and applying a threshold. The initial threshold was set to 0.95, which could eliminate effects from grey matter. However, this cropped out many important white matter voxels, particularly small structures spatially located between grey matter regions, e.g., internal and external capsules, that were vulnerable to inter-individual variabilities. We then spatially expanded the white matter mask by decreasing the threshold gradually in steps of 0.01 until the overlap between the mask and grey matter area could be visually noticed on the averaged T1 image (group mean T1 from OASIS-3). We found that 0.8 was the minimal value that could produce a clean white matter mask while retaining most of the important white matter structures. In a similar manner, a grey matter mask was also reconstructed using a lower threshold due to higher individual variabilities therein. The optimal threshold of 0.65 was specifically selected by first setting a threshold of 0.5 which then was increased gradually in steps of 0.01 until there were no overlapping voxels between white matter and grey matter. After that, the fMRI data within the white matter mask were spatially smoothed with a 4-mm full width at half maximum (FWHM) Gaussian kernel. The preprocessed results were subjected to a manual quality control procedure in which the passing criteria included: (1) all the preprocessed results must be successfully generated; (2) the maximal translations and rotations of head motion must be less than 2 mm and 2°, respectively; (3) the mean frame-wise displacement (FD) must be less than 0.5 mm^16^ and (4) the spatial normalization was acceptable by an expert’s visual inspection.

Images were acquired every second over a 154 second interval. Each image and white matter mask were reshaped into 2D (spatial coordinates × timestamps). The white matter mask was used to identify the white matter voxels in each image. A voxel level Pearson correlation coefficient was calculated across all white matter voxels in each individual image. Then, each voxel correlation coefficient was averaged across all images to create an average voxel level correlation.

### 2.2 Clustering

Clusters on the white matter voxel correlations were computed using MATLAB’s clusterdata function^17^ and linkage hierarchical clustering function^18^. The distance between each correlated voxel was measured using one minus the correlation. The computed distances were then clustered by minimizing the unweighted average between each distance.

### 2.3 Cross Fold Validation

To measure the stability of the clustering, two cross fold validations were completed. The first cross fold validation measured the stability between each scan in a session to measure the cluster repeatability. In this validation, one scan from each subject at each scan session was randomly assigned to a fold, resulting in two folds with 1,013 image scans each. The second cross fold validation measured the stability between subject populations to measure cluster reproducibility. Subjects were randomly assigned to a fold, and only the scans in the first session were used. Most subjects had two images in the first session, resulting in two folds with 355 and 356 subjects, and 708 image scans in each. Clustering of correlations between BOLD-fMRI signals at hierarchical levels between 2 and 40 was performed on each fold. The similarity between each individual cluster was measured using the Dice coefficient. The stability was measured as the mean Dice coefficients of all clusters at level.

This clustering technique was also assessed by comparing the functional clustered regions to regions identified by the JHU-DTI-SS Type-I Atlas “Eve Atlas”^19^. Since our clustering results appeared to be visually symmetric between the left and right hemisphere, we compared our identified regions to the left hemispheric regions in the Eve Atlas. In this comparison, our white matter correlations were clustered on hierarchical levels of 25 through 65. Each identified functional clustered region was compared to every Eve Atlas region using the Dice coefficient as a measure of similarity. The regions in the Eve Atlas and clustered white matter identified by the largest Dice coefficient were considered the best pair and removed from further possible pairings. This process was repeated until all regions identified in the clustered white matter were paired. The best Dice coefficients for each region were multiplied by two to create an estimate for individual cluster similarities in both hemispheres. Lastly, similarity at a given hierarchical level was measured as the mean of the best Dice coefficients at that level.

## 3. RESULTS

Clustering on the Pearson correlation coefficients across all white matter voxels successfully identified visually clear regions of the brain. On the first cross fold validation examining the scan-rescan stability, the average Dice coefficient at every hierarchical level was equal to one. On the second cross fold validation examining the population stability of clustering, the average Dice coefficient decreased as the hierarchical level increased. Dice coefficient local maximums were identified at hierarchical levels 4, 9, 13, 18, and 24, indicating possible well-made functional clusters (Figure 3).

**Figure 3.**
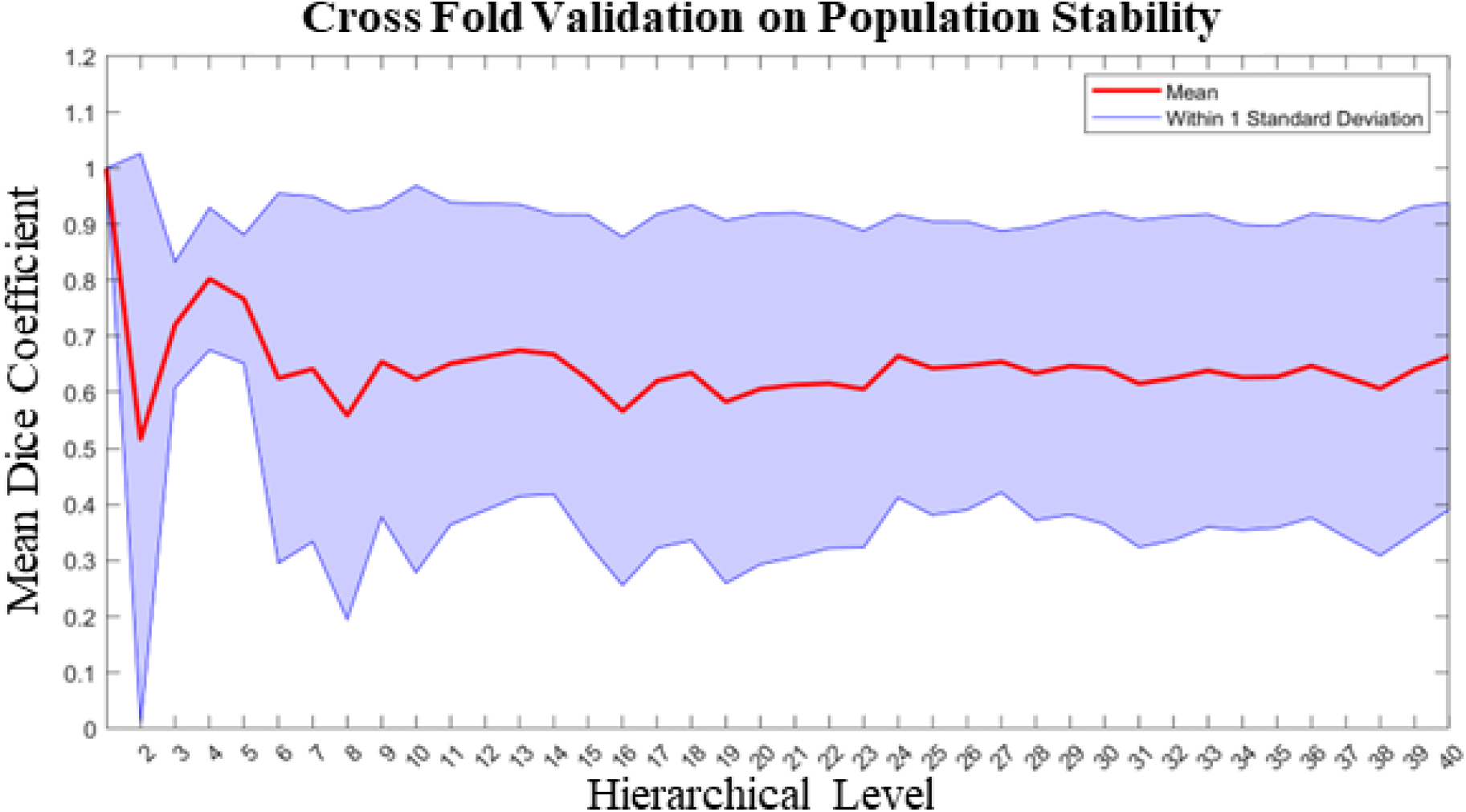
Local maximums of the average Dice coefficients occur at hierarchical levels of 4, 9, 13, 18 and 24, suggesting distinct brain regions are better identified at these levels (Figure 4). When visualizing, at a hierarchical level of 4 the major lobes are defined. As the hierarchical level increases, motor and sensory regions of the brain are identified, which are expected to be active at resting state.

### 3.1 Visualizing

Visualizing the functional clusters identified from the stability plot showed distinct white matter clusters consistent with the well-defined anatomical regions of the brain. Clustering appears symmetric across the left and right hemispheres. At all hierarchical levels, the number of clusters are equal to the hierarchical level. At lower hierarchical levels, the major lobes-occipital, parietal, coronal, and temporal-are defined. As the hierarchical level increases, motor, and sensory regions, such as the limbic lobe and motor cortex are defined (Figure 4). It is plausible for activation to be identified in these functional regions at resting state.

**Figure 4.**
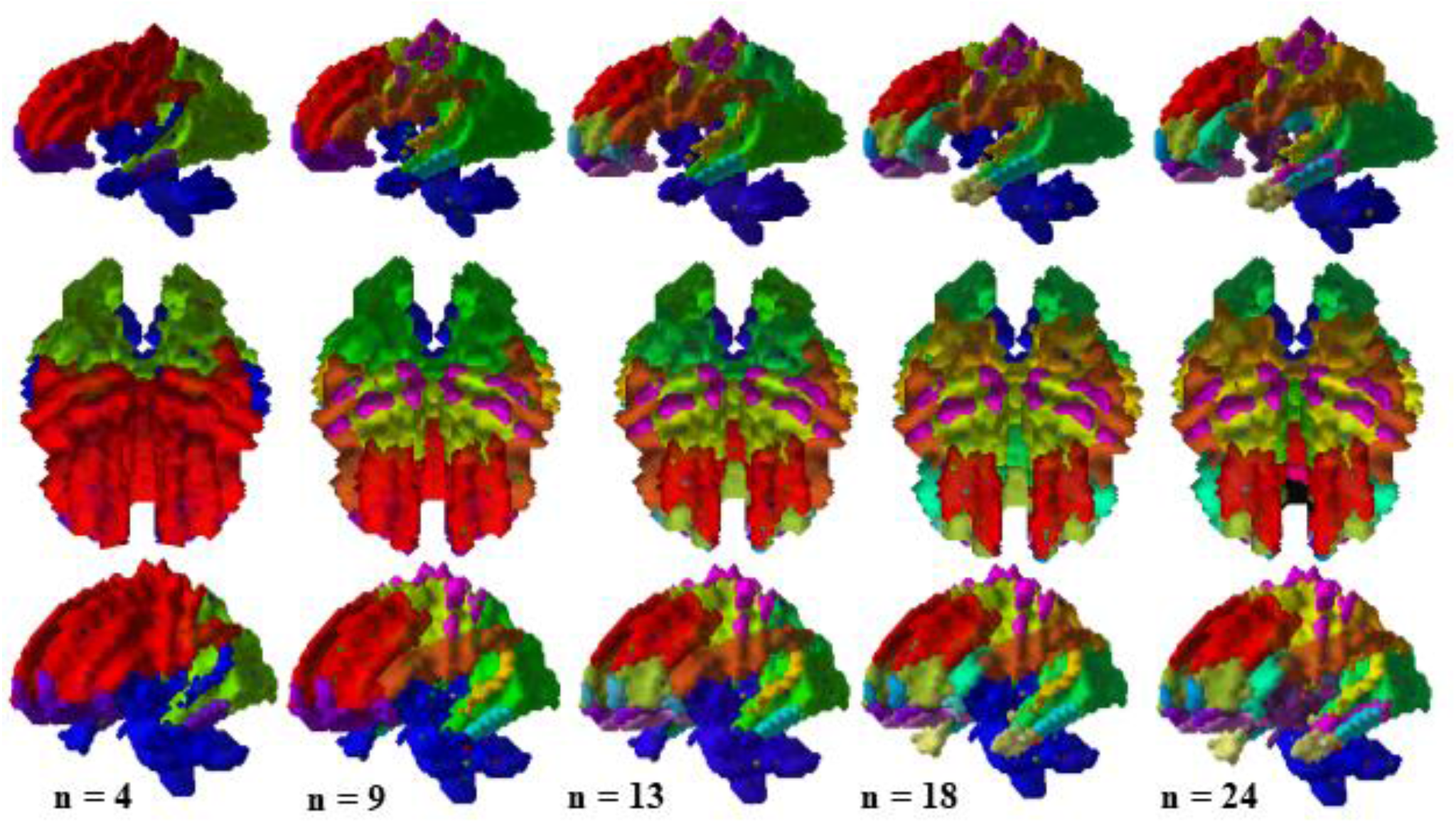
White matter clusters identified by the hierarchical clustering are consistent with well understood anatomical definitions. The number of clusters at a hierarchical level are equal to the hierarchical level. At the lower hierarchical levels (left) the regions correspond closely to the lobe regions. As the hierarchical level increases, regions are identified that are consistent with sensory and motor function regions (right).

When the functional clustered white matter regions are compared to the Eve Atlas, portions of the frontal, precentral, occipital, and orbital gyrus are consistently identified at all hierarchical levels starting at level 4. A hierarchical level of 30 is most similar to the Eve Atlas with the best mean Dice coefficient equal to 0.2881 (Figure 5). At this level, 17 of 34 frontal lobe, 2 of 10 occipital lobe, 3 of 11 parietal lobe, and 5 of 14 temporal lobe regions are identified. Portions of the cerebellum and midbrain are also identified. The primary functions of the frontal and temporal lobe are motor, memory, language, and sensory processing, which are consistent with the expected active brain regions at resting state. Within the frontal lobe, 46.05% of the corpus callosum, 37.73% of the orbital gyrus, and 41.66% of the internal capsule regions are captured by the functional clusters and all contribute to functional memory. Even at the best matched Eve Atlas regions, the Dice coefficient is much lower than the Dice coefficients determined from the stability measurement, suggesting this functional clustering is distinct from a structural parcellation.

**Figure 5.**
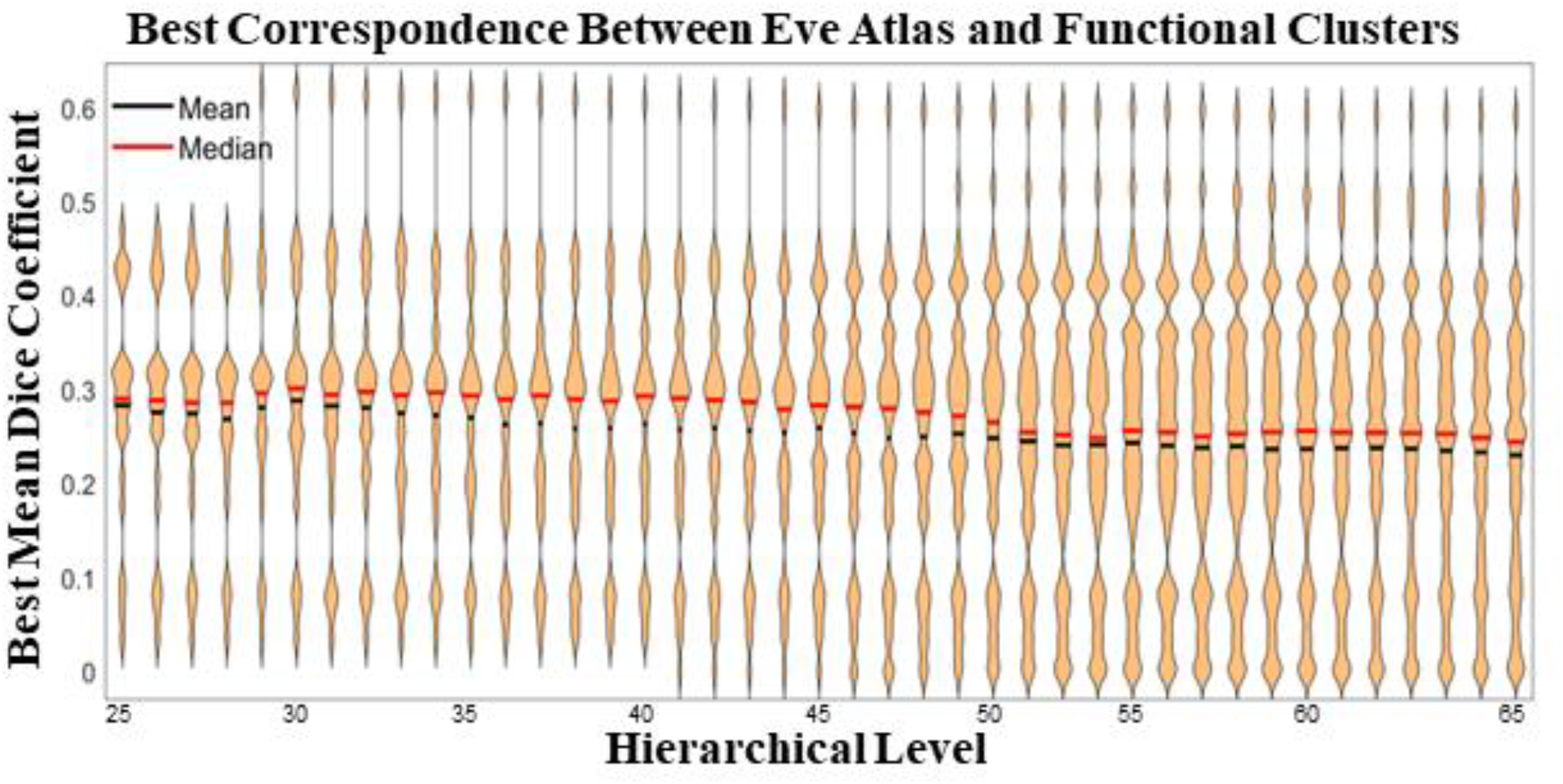
Functional clusters at hierarchical level of 30 best capture the regions in the Eve Atlas. At this level, 17 of 34 frontal lobe, 2 of 10 occipital lobe, 3 of 11 parietal lobe, and 5 of 14 temporal lobe regions are identified in the functional clusters. Although the regions are consistent with the functional memory regions that are active during resting state, the best Dice coefficients are small, suggesting functional clustering is distinct from a structural parcellation.

**Figure 6.**
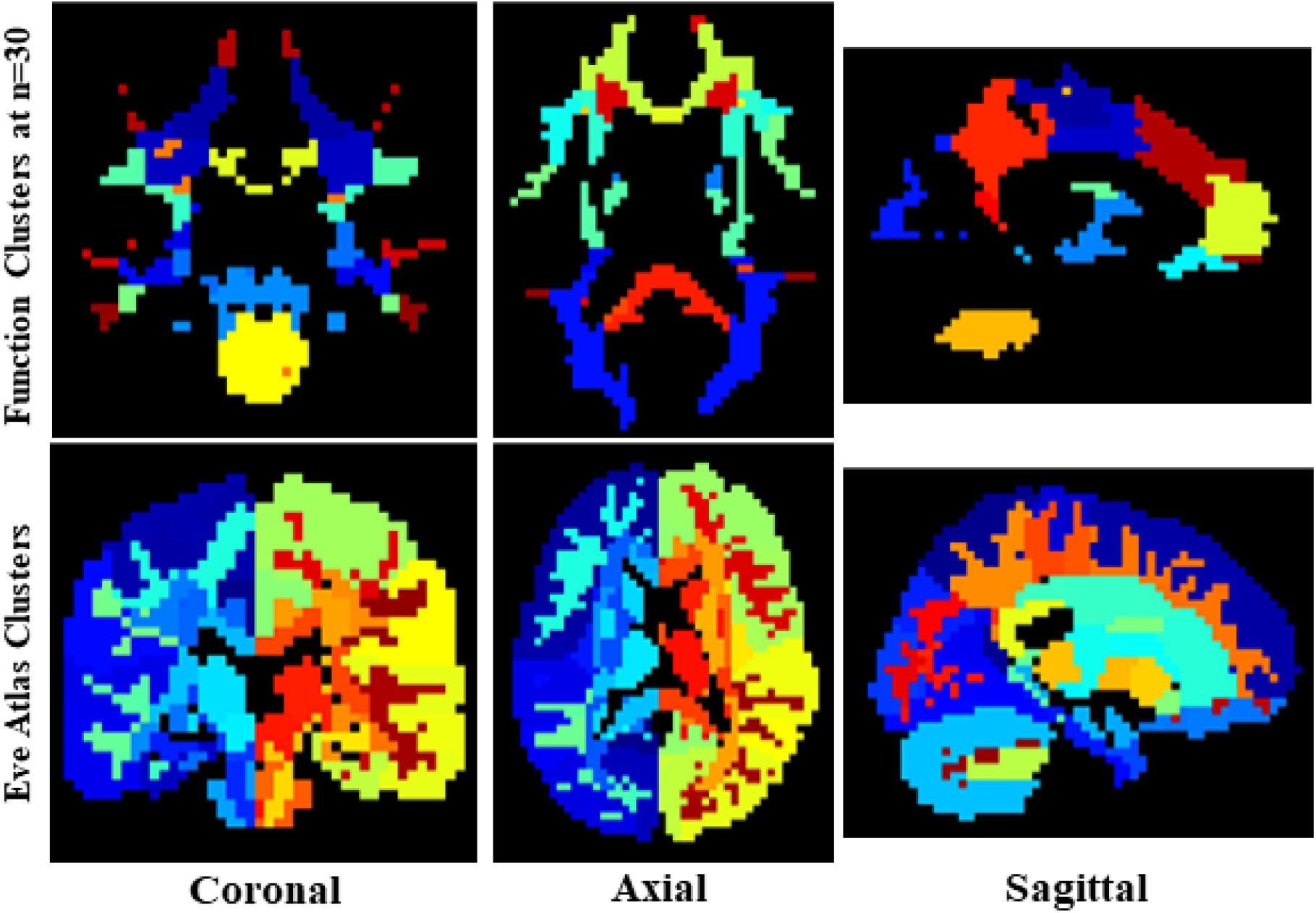
Functional clusters at a hierarchical level of 30 (top) appear highly symmetrical across hemispheres. The functional parcellation is different from the structural parcellation identified by the Eve Atlas (bottom).

## 4. DISCUSSION

Our study revealed that white matter BOLD signal is composed of functional networks, in much the same way as the more commonly analyzed gray matter BOLD signal. While there is no singular value for ‘the number of white matter networks’, we find relative stability of 6-40 clustered regions with reproducible volumetric areas in two-fold cross validation cohort. At all levels, these networks are bilaterally symmetric, with coarse levels generally representing white matter within brain lobules, and individual gyri and functionally relevant areas shown at more fine scale. Moreover, these networks do not necessarily align perfectly with well-known white matter pathways.

Our findings replicate previous studies which show white matter networks that correspond to general patterns of white matter regions and pathways^9,19,20^. However, our results suggest that these clusters, at all spatial scales, are often distinct from white matter regions. For example, average overlap coefficients between the Eve Atlas and functional clusters are generally low to moderate. Thus, there is unique information obtained from white matter functional data this is not solely derived or driven by structural connections. Future work should investigate the reliability of these networks under different image acquisition conditions, different clustering methods^9^, and different tasks.

## ACKNOWLEDGEMENTS

The project is supported by National Institutes of Health (1RF1MH123201, R01NS113832, R01EB017230, 1K01EB032898), Vanderbilt Discovery Grant (FF600670), grant of Vanderbilt Institute for Clinical and Translational Research (UL1TR0002243).

This work was supported in part by the Integrated Training in Engineering and Diabetes grant number T32 DK101003

The Vanderbilt Institute for Clinical and Translational Research (VICTR) is funded by the National Center for Advancing Translational Sciences (NCATS) Clinical Translational Science Award (CTSA) Program, Award Number 5UL1TR002243-03. The content is solely the responsibility of the authors and does not necessarily represent the official views of the NIH.

